# Tributyltin exposure leads to increased adiposity and reduced abundance of leptogenic bacteria in the zebrafish intestine

**DOI:** 10.1101/2021.07.09.451869

**Authors:** Sol Gómez de la Torre Canny, Olaf Mueller, Camil V. Craciunescu, Bruce Blumberg, John F. Rawls

## Abstract

The chemical obesogen tributyltin (TBT) is known to promote fat storage in adipose tissue through direct action on vertebrate cells. TBT also has direct toxic effects on microorganisms, raising the possibility that TBT may also promote fat storage in vertebrates by altering their microbiota. Here we show that exposure of conventionally-reared post-embryonic zebrafish to TBT results in increased adiposity, reduced body size, and altered intestinal microbiota composition including reduced relative abundance of *Plesiomonas* bacteria. To test if those microbiota alterations affected host adiposity, we exposed conventionally-reared zebrafish to intestinal bacterial strains representative of TBT-altered taxa. We found that introduction of a *Plesiomonas* strain into conventionally-reared zebrafish was sufficient to reduce adiposity and alter intestinal microbiota composition. Using new long-term gnotobiotic zebrafish husbandry methods, we found that colonization of germ-free zebrafish with *Plesiomonas* was sufficient to reduce host adiposity. Together these results show the leptogenic activity of *Plesiomonas* on zebrafish hosts, indicating that the ability of TBT to increase adiposity *in vivo* may be due in part to TBT-mediated modification of the abundance of leptogenic bacteria like *Plesiomonas*. These findings underscore how complex reciprocal interactions between animals and their microbial and chemical environments can influence energy balance and metabolic health.

**IMPORTANCE:** Obesogens are environmental chemicals that promote fat storage and are generally thought to exert this effect directly on animal cells. Using zebrafish, we show that the obesogen tributyltin can also promote fat storage by acting upon intestinal microbiota via reduction of bacteria that are sufficient to reduce fat storage.

## INTRODUCTION

Energy balance is a central aspect of animal physiology. When energy intake surpasses energy expenditure (i.e., positive energy balance), excess energy is stored as triglycerides in white adipocyte cells constituting adipose tissue (1, 2). Obesity is a condition of excessive accumulation of adipose tissue relative to body size. Obesity is a growing global public health challenge (3, 4) and a major risk factor for other non-communicable diseases (5, 6). Identification of environmental factors promoting sustained positive energy balance and expansion of adipose tissue is therefore an important goal (7). The intestinal microbiota has emerged as a key environmental factor modulating adipose tissue and body size. Mice reared germ-free (GF) in the absence of microorganisms have reduced adipose tissue (8) and body size (9) compared to mice colonized with their normal microbiota. Moreover, transplantation of fecal microbial communities into GF recipient mice from identical twins discordant for obesity (10), or from individuals with or without a form of malnutrition called kwashiorkor (11), recapitulated the respective phenotypes of the human host. Thus, both the presence and specific composition of gut microbiota have the potential to alter energy storage in adipose tissue and body size in vertebrates.

Interaction between the gut microbiota and their vertebrate hosts can be further impacted by xenobiotics (i.e., chemicals not produced by metabolisms naturally occurring in organisms and the environment) (12). Some xenobiotics can alter the composition of intestinal microbiotas and alter host physiology (13–16). Also, xenobiotics can be chemically transformed by intestinal microbiota thus modifying their toxicity, bioavailability, and efficacy when used as therapeutics (17–20).

One class of xenobiotic chemicals particularly relevant to energy balance are obesogens, which are defined by their ability to promote adipose tissue accumulation and alter metabolism (21, 22). The organotin compound tributyltin (TBT) is a prototypical obesogen. TBT was once broadly used in antifouling paints to inhibit growth of organisms on the submerged surfaces of ships, and consequently accumulated in marine sediments and organisms prior to the ban on its use (23, 24). Although TBT toxicity was first identified as causing imposex in neogastropods (25, 26), it was later shown to act as an obesogen through *in vivo* studies in multiple tetrapod species (27–31). The effects of TBT in fishes remain less understood, but include increased visceral adipose tissue in the rare minnow (*Gobiocypris rarus*) (32) and inhibited fat mobilization from adipose tissue in response to starvation in zebrafish (*Danio rerio*) (33, 34). Importantly, these *in vivo* studies were conducted in conventionally-reared (CR) animals colonized with their normal microbiota. Thus, it remains unclear whether TBT promotes adiposity via direct effects on host tissues or indirectly via their associated microbiota. Indeed, TBT directly promotes adipocyte differentiation in vertebrate cells *in vitro*, likely acting upon peroxisome proliferator-activated (PPAR) γ and retinoid X (RXR) receptors (27, 28, 35–40). The possibility that TBT might directly affect microbial physiology is supported by studies in bacteria, fungi, and other environmental microbial communities (41–49). Variable levels of microbial susceptibility to TBT have been linked to differences in TBT adsorption, bioaccumulation, enzymatic transformation, efflux and other adaptations (47, 50–58). It is therefore possible that TBT could also directly impact the composition and activity of host-associated microbial communities. Indeed, recent studies showed TBT exposure affects gut microbiota composition in rats and mice (59–61). However, the specific role of TBT-modified microbes on host adiposity remains untested, and such TBT-microbiota interactions in other vertebrates are yet unexplored.

In this study, we used the zebrafish to test if the host-associated microbiota and TBT interact to modulate host body size and adiposity. We and others have previously defined the composition of the conventional zebrafish gut microbiota (62–64) and developed short-term gnotobiotic methods for larval zebrafish to reveal impacts of specific microbiota members on host physiology (62, 65–67). We and others also showed that zebrafish develop white adipose tissues during later post-embryonic stages that display extensive homologies with those of mammals (68–71). Using TBT and microbial exposures under conventional as well as new long-term gnotobiotic husbandry we developed for this work, we tested the hypothesis that TBT affects vertebrate adiposity by modifying microbiota composition.

## RESULTS

### Tributyltin exposure affects body size and adiposity

We conducted a repeated cross-sectional study to evaluate the impact of tributyltin chloride (TBT) exposure on post-embryonic zebrafish (Fig. 1A). We raised zebrafish embryos conventionally until 18 days post-fertilization (dpf) when the intestinal microbiota had been established (64, 72). We then transferred the experimental cohort to a modified flow through aquaculture system (mFTS, see Methods), where we initiated exposure to TBT (0.01, 0.1, and 1 µg/l) or DMSO vehicle control in tank water at 22 dpf. These TBT concentrations were selected based on ranges observed in the environment and in animal tissues (73–76). Because adipose tissue development in zebrafish depends on body size, particularly on standard length (SL) (69, 70), we size-matched fish prior to exposure. We sampled fish from individual tanks at 7, 14, and 21 days post-exposure (dpE) to measure SL (77) and adiposity (defined as the ratio of the total two-dimensional area of adipose tissues per total body area excluding fins) (78), before dissecting their intestines for 16S rRNA gene sequencing analysis of the intestinal microbiota. We also sampled the tank water to evaluate concomitant effects of TBT on the aquatic microbiota (Fig. 1A, see Methods).

**Figure 1:**
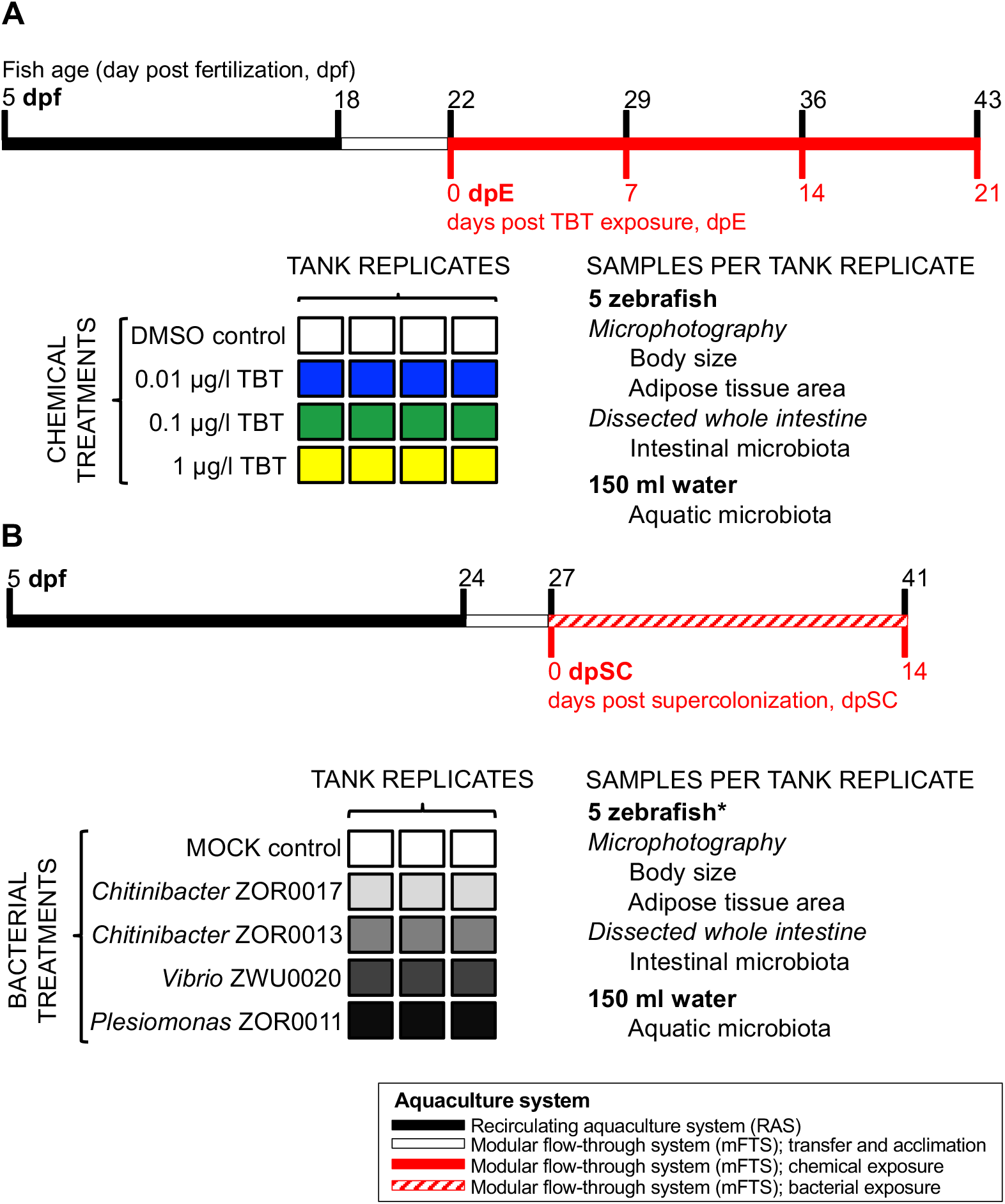
Schematic representation of *in vivo* experiments using conventionally-reared zebrafish. (A) Experimental design for exposure to tributyltin chloride (TBT). We prepared four tank replicates per chemical treatment. The concentrations of TBT in tank water are indicated in the schematic. DMSO was used as vehicle control. (B) Experimental design for supercolonization with individual bacterial strains. We prepared three tank replicates per bacterial treatment, including the four indicated bacterial strains and a mock supercolonization negative control of water from the recirculating aquaculture system (RAS). For both experiments, the timeline represents the fish age in days post-fertilization (black ticks, dpf) in relation to the days of chemical exposure (red ticks, dpE) or supercolonization (red ticks, dpSC). The different bar colors represent the aquaculture systems used for housing the zebrafish and are listed in the box below panels A and B. We sampled 5 fish per tank to measure body size and adiposity, prior to dissecting their whole intestine for 16S rRNA gene sequencing analysis of the intestinal microbial communities. We also sampled 150 ml of tank water for 16S rRNA gene sequencing analysis of the aquatic microbial communities. (*) The same sampling strategy was used in all timepoints of both experiments, except at 14 dpSC when we sampled all the remainder fish as this experiment had only two timepoints.

We detected no significant differences in zebrafish SL or adiposity between treatment groups (0 dpE; Fig. 2A and B) or replicate tanks (data not shown) prior to exposure. Although no differences in adiposity between the control and all three TBT concentrations were observed at 7 or 14 dpE, exposure to 1 µg/l TBT did cause an increase in adiposity compared to controls at 21 dpE (*P* = 0.0120; Fig. 2B and D). In addition to greater adiposity, we observed a reduction of SL in zebrafish exposed to 1 µg/l of TBT at 14 and 21 dpE (*P* = 0.0294 and *P* = 0.0105, respectively; Fig. 2A and C). Other measurements of body size including height at anterior of anal fin (HAA) (77) and total body area were only significantly reduced at 21dpE and only in the 1 µg/l TBT treatment group (*P* = 0.0042 and *P* = 0.0083; Fig. S1). Therefore, 1 µg/l TBT inhibits body growth and promotes adiposity in zebrafish after 21 days of exposure.

**Figure 2:**
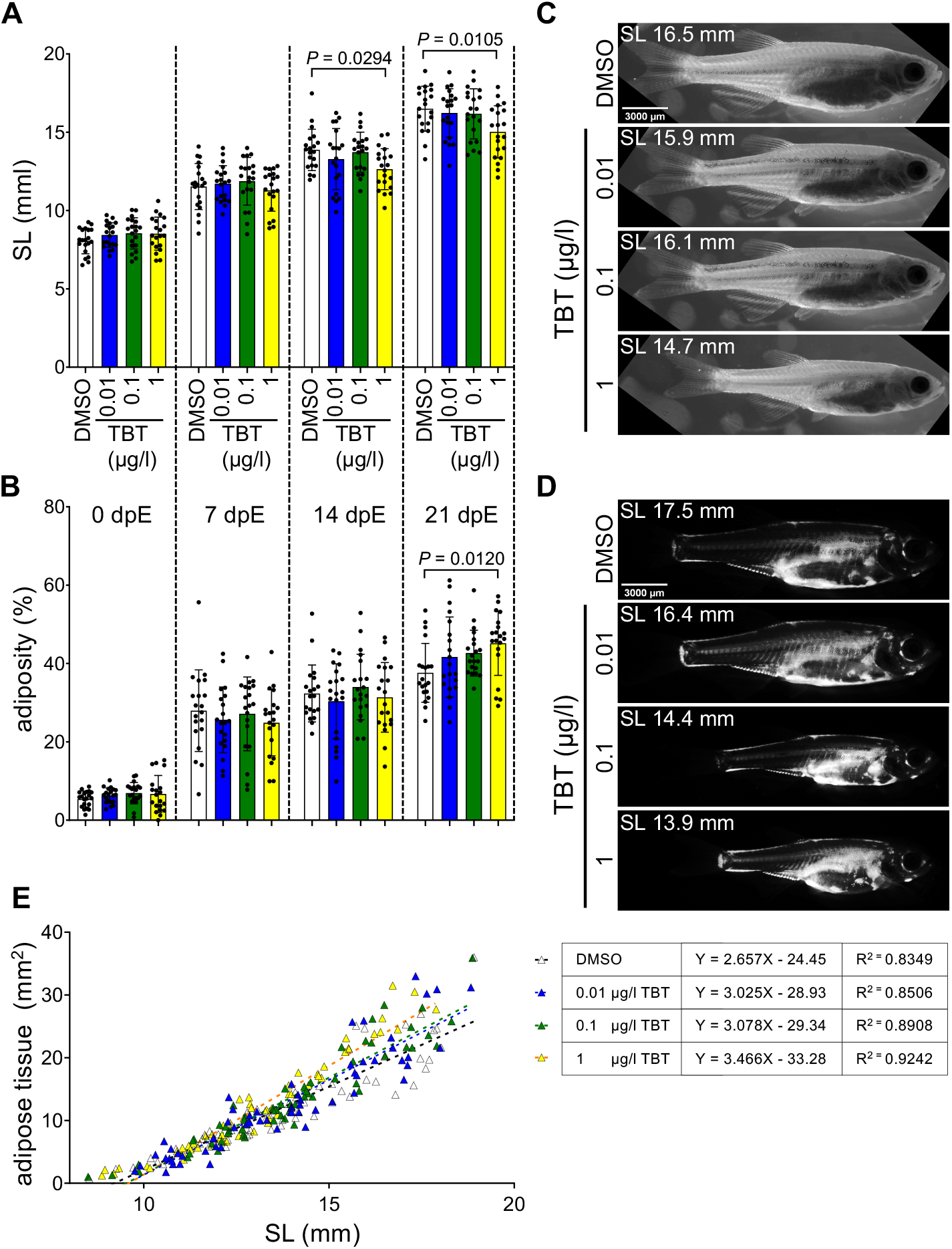
Exposure to 1 µg/l TBT increases adiposity and inhibits body size *in vivo* after 21 days. (A) Standard length (SL, the distance between the snout and the caudal peduncle) measurements. (B) Adiposity measurements (the total two-dimensional area of adipose tissues per body area excluding fins). Bars represent the mean and error bars the standard deviation. Means were compared to the DMSO control using one-way ANOVA and post-hoc Dunnett’s multiple comparison tests. When significant, the multiplicity adjusted *P*-value is reported for the comparison. (C) Brightfield images of whole fish at 21 dpE. Selected fish are representative of the mean SL per chemical treatment. (D) Nile red fluorescence images reveal adipose tissues in whole fish at 21 dpE. Selected fish are representative of the mean adiposity per treatment. Their corresponding SL is reported. (E) Linear regression analysis between SL (x-axis) and total adipose tissue area (y-axis). All samples across all timepoints (7 dpE, 14 dpE, and 21 dpE) and treatments are included. Deviation from linearity per chemical treatment was tested and found not significant (all *P* ≥ 0.0853). Equations and goodness of fit values per chemical treatment are reported on the panel. The best fit values of the slopes per treatment were compared to the DMSO control using one-way ANOVA and post-hoc Dunnett’s multiple comparison tests. The only significant adjusted *P*-value corresponded to the comparison between the DMSO and 1 µg/l TBT treatments (*P* < 0.0005).

Since zebrafish display substantial inter-individual variation in SL even when age-matched (Fig. 2A), we combined data from 7, 14, and 21 dpE to assess if TBT treatment impacted the overall developmental relationship between SL and total adipose tissue area (69, 70). A linear relationship between SL and adipose tissue area was maintained in all treatment groups (all *P* ≥ 0.0853), but the slope of the 1 µg/l TBT group was significantly greater than DMSO vehicle control (*P* = 0.0005; Figure 2E). Together, this indicates that exposure to 1 µg/l TBT alters the developmental relationship between SL and adipose tissue accumulation.

### Exposure to TBT is not a major determinant of intestinal or aquatic microbiota composition but does alter the relative abundance of specific bacterial taxa

Having observed the effects of TBT on adiposity and body size, we tested if chemical exposure also affected zebrafish intestinal or tank water microbiotas. We performed 16S rRNA gene sequencing of whole intestines and tank water sampled during TBT exposure (Fig. 1A). Intestinal and aquatic microbiota composition were significantly different from each other at all time points and TBT concentrations (all *P* ≤ 0.003; Table S1). Within each TBT concentration, the composition of intestinal microbiota at each time point was significantly different from all other time points (all *P* ≤ 0.028; Table S1). Therefore, sample type and developmental age were major determinants of intestinal microbiota composition. In contrast, TBT exposure had a relatively minor effect on intestinal and aquatic microbial community composition. We found only two instances during the study in which the composition of intestinal communities from zebrafish treated with TBT significantly differed from those in vehicle control tanks: at 7 dpE when compared to 0.1 µg/l TBT (*P* = 0.0030; Table S1) and at 14 dpE when compared to 1 µg/l TBT (*P* = 0.0430; Table S1). Notably, it was at 14 dpE that we first observed SL differences in zebrafish treated with 1 µg/L TBT. However, a significant microbiota composition difference in zebrafish treated with 1 µg/l TBT was no longer detected at the end of the experiment at 21 dpE, when both adiposity and body size phenotypes were evident (Table S1, Fig. 2, and Fig. S1). Also, we found that aquatic communities from control tanks were only significantly different compared to 1 µg/l or DMSO at 7 and 21 dpE (*P* = 0.028 and *P* = 0.029, respectively; Table S1). Comparison of indices and estimators of α-diversity between controls and all TBT concentrations revealed no differences in the total number of OTUs, Chao1, Shannon or Faith’s phylogenetic indices. Together, these data indicate that TBT does not markedly alter the overall composition or diversity of the microbial communities in zebrafish intestine or their tank water.

Although TBT exposure did not influence the overall composition of microbial communities, we reasoned that it may affect individual microbial taxa. We therefore applied linear discriminant analysis effect size analysis using LEfSe (79). LEfSe identified biomarker taxa through pairwise comparisons of the relative abundance of OTUs in microbial communities from zebrafish treated with DMSO or with 1 µg/l TBT (the only TBT concentration that rendered body size and adiposity phenotypes) in every timepoint of the exposure. We discovered 51 biomarker taxa at the genus level in the zebrafish intestine (Table S2) and 79 in water (Table S3) associated with either of the selected chemical treatments at any of the time points sampled (7, 14, and 21 dpE). Further, only 4/51 biomarkers in intestinal communities and 9/79 in aquatic communities were found in more than one of the time points sampled (Tables S2 and S3), in accord with our observation above that age is a major determinant of community composition (Table S1). Furthermore, only 7/51 biomarkers in the intestine were also identified as biomarkers in the aquatic communities, in accord with the intestinal and aquatic microbial communities being distinct (Table S1). Finally, most biomarkers in the intestine (47/51) and water (67/79) were only significantly altered at 14 or 21 dpE, the same time points when host phenotypes were also altered (Tables S2 and S3; Fig. 2). Together, these results identify bacterial genera in the zebrafish intestine and tank water that were altered by treatment with 1 µg/l TBT.

### Supercolonization of the zebrafish intestine with Plesiomonas ZOR0011 reduces adiposity but does not affect body size

We hypothesized that the effects of TBT on adiposity and body size may be mediated in part by the observed alterations in the relative abundance of specific bacterial genera. To test that possibility, we exposed CR zebrafish to specific bacterial isolates for 14 days (Figure 1B; see Methods). We reasoned that this supercolonization approach would allow us to test the effects of increasing abundance of specific community members in an otherwise normal intestinal microbial community, presumably resembling the specific changes in microbiota composition induced by TBT exposure.

We referenced LEfSe-identified intestinal biomarker taxa associated with TBT exposure and alteration in adiposity and body size phenotypes (Fig. 2; Table S2) against a collection of bacterial strains previously isolated from the zebrafish intestine (64, 80). We found strains representing biomarker genera *Plesiomonas* (Class Gammaproteobacteria, Order Enterobacterales; strain *Plesiomonas* ZOR0011), *Vibrio* (Class Gammaproteobacteria, Order Vibrionales; strain *Vibrio* ZWU0020), and *Chitinibacter* (Class Betaproteobacteria, Order Neisseriales; strains *Chitinibacter* ZOR0017 and *Chitinibacter* ZOR0013), all of which grow aerobically and were originally isolated at similar ages to that of our experimental cohort (64). In accord with our LEfSe results, the relative abundance of *Chitinibacter* OTU in the intestines of fish exposed to 1 µg/l TBT was higher compared to DMSO controls at 14 and 21 dpE (*P* = 0.0045 and *P* = 0.0224, respectively; Fig. 3A). Further, the relative abundances of *Vibrio* and *Plesiomonas* OTUs in the intestines of fish exposed to 1 µg/l TBT were lower compared to DMSO controls at 14 dpE (*P* = 0.0335 and *P* = 0.0205, respectively; Fig. 3C and E). In addition, *Plesiomonas* OTU was also more abundant in the intestines of fish treated with 0.1 µg/l TBT at 7 dpE (*P* = 0.0098; Fig. 3E). The relative abundances of these three OTUs remained unaltered in the tank water at all time points and all TBT concentrations (Fig. 3B, D, and F). These results show that TBT altered the relative abundance of all three selected bacterial genera (*Plesiomonas*, *Vibrio*, and *Chitinibacter)* in the intestine at 14 dpE, the time point of TBT exposure when the earliest host phenotype was first observed (Fig. 2), and one of the only two timepoints in which TBT was a major determinant of microbiota composition (Table S1). Based on the directionality of the TBT-induced changes in the relative abundance of these OTUs by 14 dpE, we hypothesized that an increase in *Chitinibacter* would promote adiposity and reduce body size; while an increase in *Plesiomonas* or *Vibrio* would reduce adiposity and increase body size.

**Figure 3:**
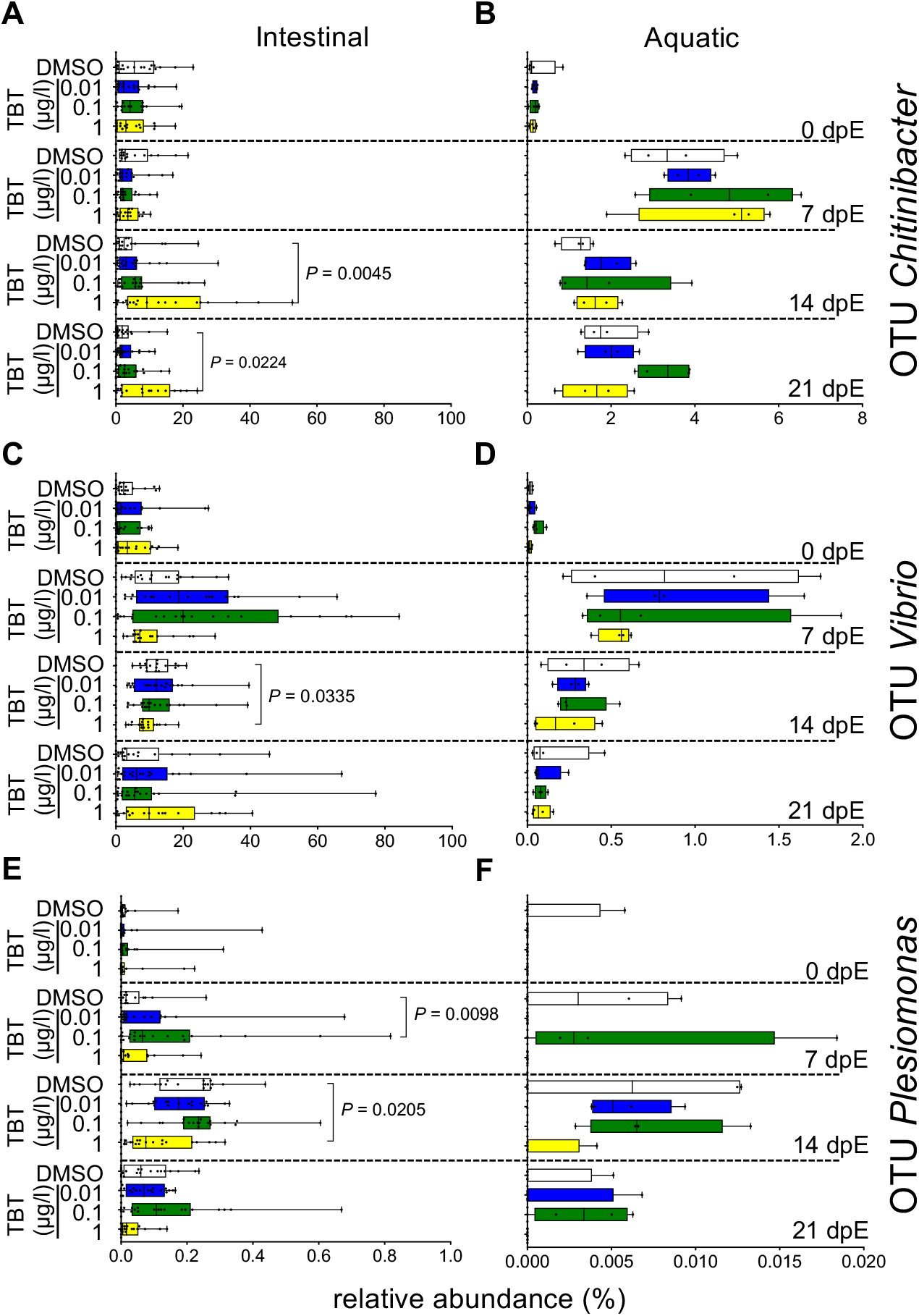
TBT exposure alters the relative abundance of specific bacterial taxa in the zebrafish intestine but not in the water. Relative abundance of selected members of intestinal microbial communities during in vivo exposure to the TBT concentrations and at the time points indicated. Selected OTUs from the intestinal (A, C, and aquatic (B, D, F) microbial communities assigned to genera *Chitinibacter* (A, B), *Vibrio* (C, D), and *Plesiomonas* (E, F). Box and whisker plots represent the interquartile range, with the line in the box defining the median. Whiskers represent the minimum and maximum value. The mean of each bacterial treatment was compared to the control using the Mann-Whitney test per time point and sample types. When significant, the *P*-values are reported for the comparisons.

Zebrafish entering the supercolonization experiment were size-matched to those that had entered the *in vivo* TBT exposure experiment to ensure they were at similar developmental stages. Mock supercolonization using only autoclaved RAS water was used as a negative control. We sampled fish from individual tanks to evaluate the impact of the bacterial treatments on body size, adiposity, as well as intestinal and aquatic microbial communities (Fig. 1B). Prior to exposure, measurements of zebrafish adiposity and body size did not reveal differences between experimental groups (Fig. 4A and B). However, after 14 days of supercolonization, animals treated with *Plesiomonas* ZOR0011 had a significant reduction in adiposity (*P* = 0.0334; Fig. 4B and without effects on standard length or other body size measurements (Fig. 4A and C; Fig. S1). The other three bacterial strains did not significantly affect either of these traits. The observation of the predicted adiposity phenotype after supercolonization with *Plesiomonas* ZOR0011 supports the possibility that the reduction in the relative abundance of *Plesiomonas* bacteria following TBT exposure may mediate, at least partially, the resulting host phenotypes.

**Figure 4:**
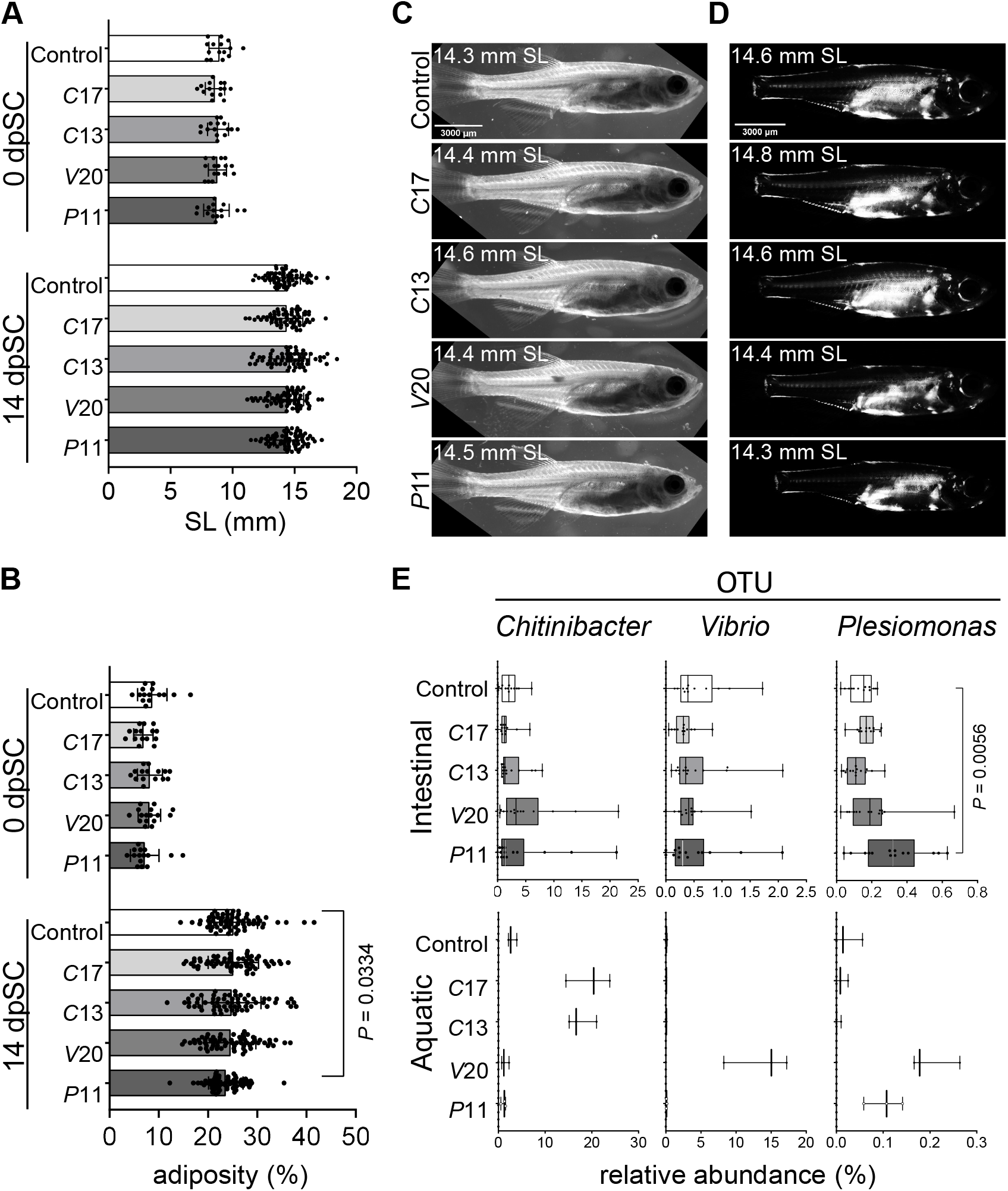
Supercolonization with *Plesiomonas* ZOR0011 is sufficient to inhibit adiposity. Zebrafish were supercolonized daily for 14 days with one of four bacterial strains: *Chitinibacter* ZOR0017, C17; *Chitinibacter* ZOR0013, C13; *Vibrio* ZWU0020, V20; or *Plesiomonas* ZOR0011, P11. (A) Standard length (SL) and (B) adiposity measurements before initiating the bacterial treatment (0 days) and after 14 days of daily supercolonization (dpSC). Bars represent the mean and error bars the standard deviation. Means were compared to the negative control using unpaired t-test. When significant, the *P*-value is reported for the comparison. (C) Brightfield images of whole fish representative of the mean SL per treatment at 14 dpSC. (D) Nile red fluorescence images reveal adipose tissues in whole fish representative of the mean adiposity per treatment at 14 dpSC. Their corresponding SL is reported. (E) Relative abundances of OTUs assigned to genera *Plesiomonas*, *Chitinibacter*, and *Vibrio* in aquatic and intestinal microbial communities at 14 dpSC. Box and whisker plots represent the interquartile range, with the line in the box defining the median. Whiskers represent the minimum and maximum value. The mean of each bacterial treatment was compared to the control using the Mann-Whitney test. When significant, the *P*-values are reported for the comparisons.

We next used 16S rRNA gene sequencing to verify if supercolonization effectively increased the relative abundance of the introduced bacterial strains in the water or zebrafish intestinal microbial communities. We examined the relative abundance of *Chitinibacter*, *Plesiomonas*, and *Vibrio* OTUs in the aquatic communities after 14 days of supercolonization with the four different intestinal bacterial strains. None of those bacterial treatments significantly increased the relative abundance of *Chitinibacter*, *Plesiomonas* or *Vibrio* OTUs in the tank water when compared to negative controls (Fig. 4E). Nevertheless, results from the more sensitive LEfSe analysis revealed that the OTUs assigned to each introduced bacterial strain were in fact biomarker genera of their corresponding treatment group (Table S4). Therefore, supercolonization effectively increased the relative abundance of introduced bacterial genera in the aquatic environment, thereby potentially allowing them to also colonize the zebrafish intestine. However, changes in the relative abundance of members of the intestinal microbial communities of supercolonized animals were unique to the strain *Plesiomonas* ZOR0011. The relative abundance of OTU *Plesiomonas* in the intestine was increased upon supercolonization with *Plesiomonas* ZOR0011 (*P* = 0.0022; Fig. 4E), a result confirmed by LEfSe analysis (Table S5). For the other three strains tested, the respective OTUs were not altered nor identified as biomarkers by LEfSe analysis. These data support the hypothesis that the host adiposity phenotype caused by *in vivo* exposure to 1 µg/l TBT may be mediated in part by the chemically-induced alteration of the relative abundance of *Plesiomonas* in the zebrafish intestine.

### Supercolonization with Plesiomonas ZOR0011 alters zebrafish intestinal microbiota

We next asked if supercolonization with *Plesiomonas* ZOR0011 or any of the other selected bacterial strains, altered the composition of the intestinal or aquatic microbiota. As expected, composition of intestinal and aquatic microbiotas remained significantly different regardless of the treatment (all *P* ≤ 0.008; Fig. S2; Table S6). Supercolonization with the different bacterial strains did not alter α-diversity of aquatic or intestinal communities (total number of OTUs, Chao1, Shannon or Faith’s phylogenetic indices). However, the composition of intestinal microbial communities in zebrafish supercolonized with *Plesiomonas* ZOR0011 or *Chitinibacter* ZOR0017 displayed significant differences compared to negative controls (*P =* 0.001 and *P* = 0.027 respectively; Fig. S2, Table S6). Also, the intestinal microbial communities in zebrafish supercolonized with *Plesiomonas* ZOR0011 and *Chitinibacter* ZOR0017 are distinct (*P =* 0.001; Fig. S2, Tables S6). These results show that supercolonization with *Plesiomonas* ZOR0011 and *Chitinibacter* ZOR0017 had distinct effects on intestinal microbiota composition.

### Monoassociation of zebrafish with Plesiomonas ZOR0011 inhibits adiposity and body size

Supercolonization with *Plesiomonas* ZOR0011 reduced adiposity, but also altered the composition of the established intestinal microbiota. We therefore sought to test the sufficiency of *Plesiomonas* ZOR0011 to directly influence adiposity and body size in the absence of other microbes using a gnotobiotic approach. Although methods for gnotobiotic husbandry of zebrafish embryos and larvae have been established, they were not optimized to support growth of zebrafish into later stages when adipose tissues develop (68–70). We therefore developed new methods for long-term gnotobiotic zebrafish husbandry (see Methods) to raise germ-free (GF) zebrafish to 14 dpf before conducting a monoassociation with *Plesiomonas* ZOR0011 or *Chitinibacter* ZOR0013. *Chitinibacter* ZOR0013 was selected as a control because it did not elicit a phenotype nor altered the composition of the intestinal microbiota after supercolonization. As an additional reference control, we tested the impact of TBT on zebrafish body size and adiposity in the absence of any microbes by exposing GF zebrafish to 1 µg/l TBT or DMSO. We subjected GF zebrafish to these treatments for 21 days to match the length of adiposity phenotype during the TBT exposure (Fig. 4B and D). We detected no differences in mortality (all *P* ≥ 0.1986 at 7 dpE/dpC; and *P* ≥ 0.5213 at 21 dpE/dpC) or the density (all *P* ≥ 0.8869 at 7 dpE/dpC; and *P* ≥ 0.9474 at 21 dpE/dpC) of the flasks between the chemical or bacterial conditions tested (Methods). Whereas there were no phenotypical alterations in fishes after 7 days of treatment, we observed a reduction in adiposity of gnotobiotic zebrafish 21 days post-colonization (dpC) with *Plesiomonas* ZOR0011 (*P* = 0.0399; Fig. 5A, B, and D), resembling the effect of supercolonization of CR zebrafish with this bacterial strain (Fig. 4B and D). In contrast to its supercolonization phenotype, monoassociation with *Plesiomonas* ZOR0011 also reduced body size of gnotobiotic zebrafish (*P* = 0.0256; Fig. 5A and C). Finally, neither monoassociation with *Chitinibacter* ZOR0013 or exposure to 1 µg/l TBT in the GF background elicited body size or adiposity phenotypes after 21 days (Fig. 5). Together, these results show that *Plesiomonas* ZOR0011 is sufficient to directly reduce adiposity and body size, and suggest that obesogenic effects of TBT on zebrafish may require the presence of microbiota.

**Figure 5:**
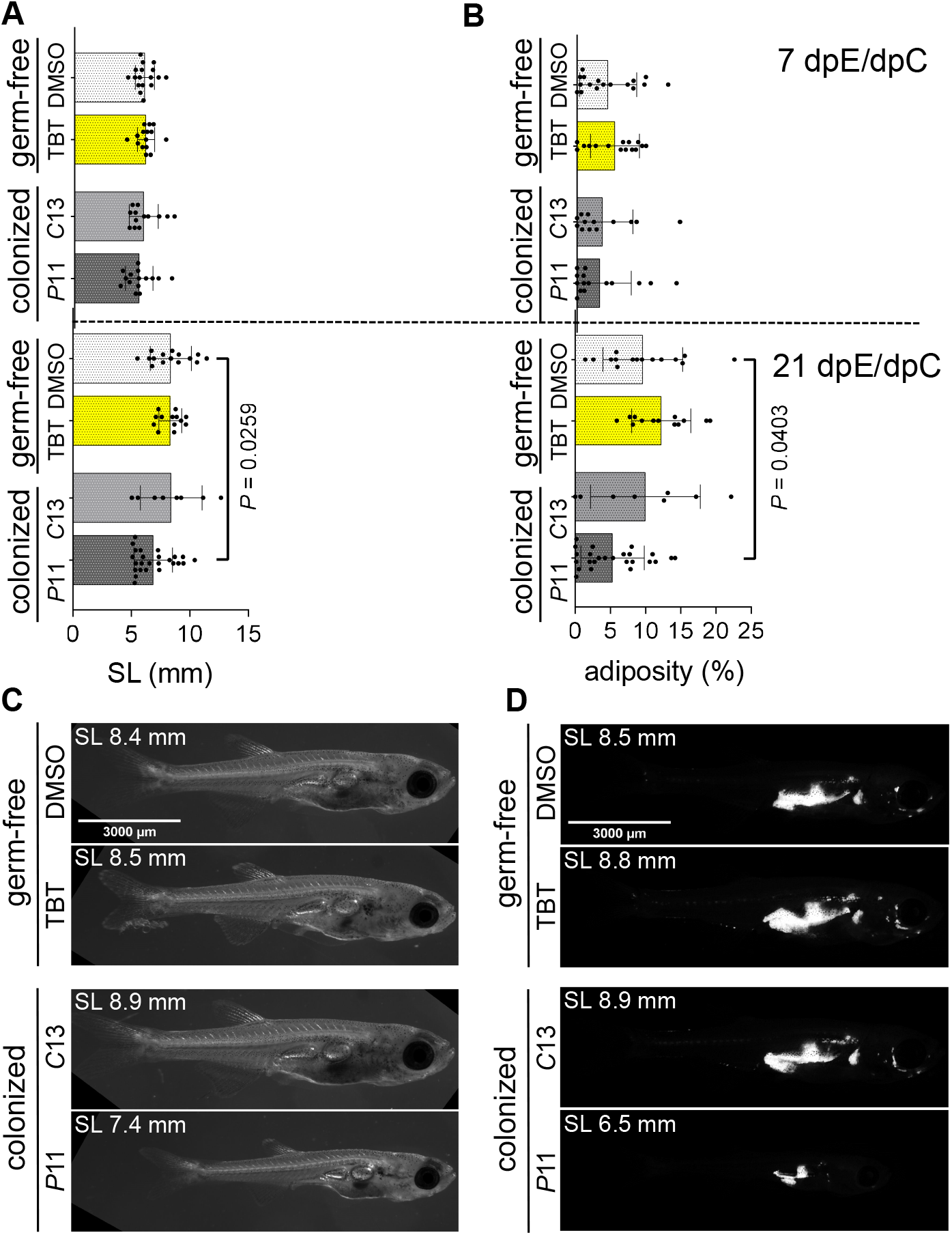
Monoassociation reveals that *Plesiomonas* ZOR0011 is sufficient to reduce body size and adiposity in zebrafish without an established microbiota. (A) Standard length and (B) adiposity after exposure of germ-free zebrafish to TBT; or monoassociation (colonized) with *Chitinibacter* ZOR0013 (*C*13) and *Plesiomonas* ZOR0011 (*P*11). Measurements were taken 7 and 14 days post-exposure (dpE) or post-colonization (dpC). Bars represent the mean and error bars the standard deviation. Means of each treatment were compared to the DMSO control using one-way ANOVA and the post-hoc Dunnett’s multiple comparison test. When significant, the multiplicity adjusted *P*-value is reported for the comparison. (C) Brightfield images of whole fish representative of the mean SL per treatment measured after 14 days. (D) Nile red fluorescence images reveals adipose tissues in whole fish representative of the mean adiposity per treatment measured after 14 days. Their corresponding SL are reported.

### TBT affects specific members of the intestinal microbiota directly in the absence of the zebrafish host

The studies above did not resolve whether TBT affects *Plesiomonas* and other intestinal bacteria *in vivo* directly, or indirectly through host-dependent processes. To investigate the direct effect of TBT on zebrafish intestinal bacteria, we harvested intestinal microbiota from CR zebrafish and cultured them *in vitro* in non-selective rich media containing different concentrations of TBT similar to those tested in the *in vivo* chemical exposure (0.01, 0.1, and 1 µg/l TBT) and two additional higher concentrations (10 and 1000 µg/l TBT) anticipating the presence of highly-resistant bacteria observed in some aquatic environments (42). Zebrafish used as microbiota donors were size-matched to zebrafish from control tanks at the endpoint of the *in vivo* TBT exposure (see Methods). We incubated the communities *in vitro* for 24 hours at room temperature under aerobic and anaerobic conditions and used 16S rRNA gene sequencing to interrogate their composition. Of a total of 272 OTUs detected across the different treatments and the input community, only 62 were observed both in the input community and at least one condition after *in vitro exposure to TBT*. Nevertheless, most of the OTUs detected in the *in vitro* cultures (85/90) were also observed in the intestinal microbiota of zebrafish at 21 dpE during the *in vivo* chemical exposure in at least one of the chemical conditions. Therefore, although OTUs from the *in vitro* experiments represent only the culturable fraction of an *in vivo* community, they provide a relatively broad representation of OTUs found in the intestinal microbiota of zebrafish from the same size that underwent TBT exposure *in vivo* (Table S7).

As in the *in vivo* chemical exposures, diversity analysis showed that TBT was not a major determinant of the composition of the culturable zebrafish intestinal microbiota at any of the concentrations tested *in vitro* (all *P* ≥ 0.05, Table rename). In addition, α-diversity was not altered by the exposure to TBT (total number of OTUs, Chao1, Shannon or Faith’s phylogenetic indices). LEfSe analysis identified 37 biomarker taxa of *in vitro* TBT exposure at the genus level, in at least one of the pairwise comparisons between the DMSO and the different TBT concentrations (Table S8). Of the 37 biomarker genera, 11 of them were also identified as biomarkers of the *in vivo* exposure to 1 µg/l TBT at time points when zebrafish phenotypes were detected (14 and 21 dpE, Table S8). These 11 biomarkers included genera *Vibrio and Plesiomonas*. *Plesiomonas* showed an increased relative abundance (2.7-fold) only at the highest TBT concentration of 1000 µg/l under aerobic conditions. In contrast, the lower TBT concentrations used in the *in vivo* chemical exposure experiment (0.01, 0.1, and 1 µg/l TBT) did not affect the relative abundance of *Plesiomonas* in microbiota cultured *in vitro* (Tables S7 and S8). These results suggest that *Plesiomonas* sensitivity to TBT may be determined in part by the host.

## DISCUSSION

TBT is considered a prototypical obesogen (27–31). In zebrafish and other teleosts, TBT promotes the accumulation of lipids in different tissues and organs (32, 81–88). Exposure of post-embryonic zebrafish to TBT impaired adipose tissue mobilization in response to food restriction (33, 34), but its impact on adipose tissue accumulation in fishes was unknown. Our results establish that a 21-day TBT exposure promotes adiposity in zebrafish, indicating its obesogenic activity on teleost fishes. We also found that the TBT-induced adiposity phenotype was accompanied by reduced body size. This is in accord with reports that TBT compounds reduced body length in zebrafish (89), other teleost species (90, 91) and in *Xenopus tropicalis* (92), as well as body weight in other vertebrates (93–95), although these studies did not examine co-occurring lipid or adipose phenotypes. In humans, stunted growth and other forms of malnutrition coexist with obesity (96) and earlier manifestations of undernutrition may predispose to obesity (97). We observed that the impacts of TBT on SL (by 14 dpE) preceded overt impacts on overall body size and adiposity (by 21 dpE), but it remains unclear if these adiposity and body size phenotypes are mechanistically linked. It is possible that TBT-induced impairments in fat mobilization from adipose tissues in zebrafish (34) reduce the energy available for growth. Identification of the direct and indirect mechanisms by which TBT affects the relationship between adiposity and body size is therefore warranted.

There is a growing appreciation that xenobiotics may exert their impacts on animal health by altering the composition of their microbiotas (15, 16, 59–61, 98–100). While TBT has well-documented direct actions on adipocyte differentiation and other aspects of host physiology (27, 28, 35–40), it was only recently shown to alter the composition of gut microbiota in rodents (59–61). In attempt to test if TBT-modified gut microbiota is sufficient to affect host physiology, Zhan and coworkers transplanted TBT-modified fecal microbiota into recipient mice pre-treated with antibiotics to deplete their resident microbiota, and recapitulated adiposity and metabolic phenotypes observed in donors (60). However, this experimental approach does not control for potential confounding effects of antibiotic-resistant microbes, the direct effects of antibiotics on the recipient host (101), or the possibility of bioaccumulation of TBT in fecal microbiota. We found that TBT altered the relative abundance of a subset of intestinal bacterial genera in CR zebrafish. Whereas the alterations in intestinal microbiota composition was relatively modest, this information allowed us to identify TBT-affected bacterial genera for further sufficiency tests. It also remains possible that, beyond these compositional changes, TBT affects the functional potential of the microbiota as has been demonstrated with other xenobiotics (102–104).

We developed multiple experimental approaches to test if microbes affected by TBT exposure *in vivo* were themselves sufficient to evoke the phenotypes observed in their hosts. Supercolonization is a strategy that introduces microbial strains into animals already colonized with a microbiota, and has been effective for defining the impact of the introduced microbes on the host and the intestinal ecosystem (105–107). By supercolonizing CR zebrafish with available individual bacterial strains representing TBT-affected genera, we found that *Plesiomonas* ZOR0011 was the only strain, amongst those tested, sufficient to reduce adiposity in zebrafish. One advantage of the supercolonization approach is that it inherently accommodates the potential for microbial interactions. Indeed, *Plesiomonas* ZOR0011 supercolonization altered the microbiota composition, making it unclear whether the impact of *Plesiomonas* ZOR0011 supercolonization on host adiposity was direct or indirect via alterations in the abundance of other microbiota members. For these reasons we used a complementary gnotobiotic approach to study the isolated effect of *Plesiomonas* ZOR0011.

Although methods for gnotobiotic zebrafish husbandry have been in use for over 17 years (62, 67, 108), almost all studies have focused on early free-feeding larval stages. This postponed the need to develop nutrition and housing systems for raising zebrafish into later life stages. The husbandry methods that we developed for this study permit growth of zebrafish to standard lengths at which most adipose tissue depots develop (70). We anticipate these methods will also be useful for testing the role of microbial exposures on other aspects of post-embryonic physiology in the zebrafish. Using these new methods, we showed that the exposure of GF zebrafish to a TBT concentration that was phenotypical in CR zebrafish did not affect body size or adiposity. This suggests that GF zebrafish may be resistant to TBT-induced phenotypes, but additional studies are needed to determine if this is limited to specific developmental stages or TBT dosages. Nevertheless, our combined approaches showed that *Plesiomonas* ZOR0011 was sufficient to inhibit adiposity in zebrafish, both in the absence and the presence of a conventional microbiota.

A major finding of this study is the discovery that a bacterial strain isolated from the zebrafish intestine, *Plesiomonas* ZOR0011, has leptogenic effects on zebrafish hosts. *Plesiomonas* bacteria are common members of the intestinal microbiota in laboratory-reared zebrafish at all life stages and across multiple aquaculture facilities, and have also been observed in adult zebrafish recently captured from their natural habitat (63, 64). *Plesiomonas* abundance in the zebrafish intestine has also been shown to be sensitive to other xenobiotics besides TBT (109–113), but this study is the first to explore its direct physiologic effects on this host. Although the relative abundance of *Plesiomonas* in the intestine was relatively low in our study, it had a significant effect on host physiology and on microbiota composition. Other studies have also underscored how members of the microbiota of relatively low abundance can have significant effects on host-microbe commensalism (66, 114). The mechanisms by which *Plesiomonas* ZOR0011 exerts its leptogenic effects on zebrafish and affects microbiota composition remain unknown. The genus *Plesiomonas,* thought to consist of a single species *P. shigelloides,* is common in aquatic ecosystems and inhabits diverse aquatic vertebrates and invertebrates (115). *Plesiomonas* has also been detected in the intestinal microbiota in healthy humans and some mammals, yet also frequently associated with gastroenteritis in humans (115). We did not observe any overt pathology or mortality in zebrafish supercolonized or monoassociated with *Plesiomonas* ZOR0011, thus its sufficiency to reduce adiposity and body size may be consistent with a role on digestive physiology or energy balance. Moreover, our recent finding that the relative abundance of *Plesiomonas* members was markedly reduced by starvation in adult zebrafish, suggests potential roles in host nutritional physiology (116). Further studies are needed to discern the mechanisms by which *Plesiomonas* ZOR0011 exerts its leptogenic effect.

Whereas the focus of this study was to assess impacts of TBT on gut microbiota and its consequences *in vivo*, we also investigated the direct effect of TBT on zebrafish microbiota *in vitro* in the absence of the host. Our results demonstrated that the impact of xenobiotics like TBT on members of the zebrafish intestinal microbiota can differ between *in vivo* and *in vitro* scenarios. Many genera including *Plesiomonas* were not affected by phenotypical levels of TBT *in vivo* (0.1 and 1 µg/l TBT), and their relative abundance *in vitro* was only significantly affected by much higher concentrations (1000 µg/l TBT). We were unable to test the effect of TBT exposure to ≥ 10 µg/l on intestinal microbiota *in vivo* as this concentration caused >92% mortality as early as 1 dpE (see Methods). The global increase in synthetic chemicals like TBT is recent in the shared evolutionary history of vertebrate animals and their commensal bacteria with whom they continue to co-evolve (117–119). The effect of chemical novelty on intestinal bacteria like *Plesiomonas* is presumably an anthropogenic impact on organisms across the tree of life, as individuals or as holobionts. Zebrafish, a well-established animal model for human disease, can also serve as a model for wild and farmed fishes, vulnerable to xenobiotic exposure and central to food security (120–122). Further understanding of the specific effect of xenobiotics on intestinal microbiota can contribute to both planetary sustainability and the improvement of human and animal health.

## MATERIALS AND METHODS

### Conventional zebrafish husbandry and obtention of experimental cohorts

All zebrafish experiments were approved by the Institutional Animal Care and Use Committees of Duke University Medical Center under protocol A115-16-05 in accordance with the Public Health Service Policy on the Human Care and Use of Laboratory Animals under the United States of America National Institutes of Health Office of Laboratory Animal Welfare (OLAW). Adult zebrafish (wild-type Ekkwill strain) were raised under conventional conditions on a Pentair recirculating aquaculture system (RAS) and set up for natural breeding to obtain the embryos constituting the different experimental cohorts. Four adult fish (1:1 male to female ratio) from five families were set in mating boxes for group spawning. The resulting clutches of embryos were collected synchronously, transferred to Petri dishes containing RAS water, and maintained at a density of ∼70 fish per dish and incubated at 28 °C. Abnormal embryos were removed during the first 24 hours post-fertilization (hpf). At ∼36 hpf, embryos were pooled and randomized into groups of 62 embryos which were transferred to clean 3-liter tanks (30 tanks total). Each 3-liter tank, filled with 500 ml of RAS water, was kept static until 5 dpf. At 5 dpf, the 3-liter tanks were connected to the RAS and maintained according to our standard husbandry protocols (123) with one modification to feeding before their transfer to the modular flow through aquaculture system (mFTS, see below) where the experimental cohorts were housed during the experiments. The modification was that, instead of receiving an exclusive powdered diet of Zeigler AP100 <50-micron larval diet (Pentair, LD50-AQ) twice daily from 5 dpf until 14 dpf, larvae were supplemented twice daily with 500 µl of *Artemia* (Brine Shrimp Direct, BSEACASE) that was collected, rinsed and resuspended in an equal volume of RAS water.

The experimental cohort for the *in vivo* chemical exposure to TBT was transferred to the mFTS on 18 dpf. On the day of transfer to the mFTS, zebrafish were collected from the RAS in sterile 25 mm-deep Petri dishes, where they were anesthetized using 0.2 g/L buffered Tricaine to measure their SL (77) for size-matching individuals between 7 and 11 mm. The SL of the experimental cohort prior to TBT exposure (0 days post-exposure, dpE) was of 8.39 ± 0.82 mm (mean ± SD), which was measured at 22 dpf. A total of 20 pools of 30 individuals constituted the experimental cohort during the *in vivo* chemical exposure study. For the supercolonization experiments, the cohort was obtained in the same manner with the exception that transfer to the mFTS occurred on 24 dpf. Zebrafish entering the supercolonization experiment were size matched to those that had entered the TBT exposure to avoid developmental differences between these two experiments. The SL of the experimental cohort prior to bacterial supercolonization (0 days post-supercolonization, dpSC) was of 8.77 ± 0.84 (mean ± SD), which was measured at 27 dpf. 15 pools of 30 individuals constituted the experimental cohort during the supercolonization study.

### Modular flow through system (mFTS) construction and operation

Because TBT is likely degraded by filtration through activated carbon (124) and UV exposure (125) which are used to maintain water quality on the RAS, we designed, built and installed a modular flow-through system (mFTS) in one of the racks at our RAS. The mFTS housed the experimental cohorts for both the *in vivo* TBT exposure and the supercolonization experiments.

Each shelf in our facility fits five 10-liter polycarbonate tanks connected to the RAS by five manifolds that deliver pre-conditioned RAS water into the experimental tanks. The mFTS used the standard polycarbonate tanks from the RAS system which we modified by drilling and installing polycarbonate food-grade beverage dispenser faucets serving as spigots for draining the tanks during water exchanges (2712212, Carlisle). Inside the tanks, the spigots were covered with fiberglass mesh (33105, Lowe’s) serving as baffles to prevent fish from escaping during water exchanges. The spigots from the five tanks in each shelf were connected through fitted vinyl hoses to a 3/4-inch PVC pipe mounted with a slight angle of inclination with respect to the fish tank shelf. This angle prevented backflow during the water exchanges. One PVC pipe was used to collect the outflow of 5 tanks from a shelf. Four PVC-pipes were finally connected, through a 90-degree 3/4-inch elbow, to a fitted 1-inch vinyl hose and secured with an adjustable clamp (57188, Lowe’s). The 1-inch vinyl hose ran perpendicular to the fish tank shelves into a drain, to collect the outflow of the mFTS system. Thus, the mFTS thus allowed us to drain several fish tanks simultaneously to perform water exchange in a relatively automated and uniform manner. The outflow of each fish tank was controlled by the location of the spigot along the height of the tank. We installed the spigot to regulate an outflow of 5.5 L, with 2.5 L retained below the spigot after draining. After draining, the tanks in the mFTS were refilled with the conditioned water from the RAS system via the RAS manifold. The modified tanks in the mFTS were initially filled with 7.5 L of conditioned RAS. A total of 1.6 volumes of water were exchanged daily to maintain an adequate water quality.

The mFTS held twenty 10-liter modified polycarbonate tanks during the *in vivo* chemical exposure to TBT, and fifteen during the supercolonization experiments. Water quality parameters were assessed in the mFTS during the two studies. Conductivity was measured in the conditioned RAS water before water exchange. pH, nitrate and nitrite, and toxic ammonia were measured in the mFTS and in the conditioned water using test kits according to the manufacturer’s instructions (147008, 2745425, and 224100 respectively, HACH). Water quality parameters were measured on every tank in the mFTS on the day before adding the experimental cohort to the tanks, and on every time point sampled during the studies. The pH and conductivity in the conditioned water used for the water exchange was recorded daily. The nitrite levels in the conditioned water were 0 ppm, nitrate between 2 and 10 ppm, and toxic ammonia was 0 mg/l. During the *in vivo* chemical exposure to TBT, the pH in the tanks ranged between 7 and 7.6, nitrite between 0 and 0.15 ppm, nitrate between 2 and 10 ppm, and toxic ammonia between 0 and 0.00521 mg/l. The water pH and nitrate in the experimental tanks during the chemical exposure was comparable to that of the conditions water, while the nitrite and the toxic ammonia were higher. During the supercolonization experiments, water pH in the experimental tanks ranged between 7.37 and 7.41, nitrite was 0, nitrate ranged between 2 and 10 ppm, and toxic ammonia between 0 and 0.00106 mg/l. When compared to the conditioned water, the pH was lower, the nitrite and nitrate were comparable, and the toxic ammonia was slightly higher. Finally, the conductivity in the RAS water used during the water exchanges during the *in vivo* exposure to TBT ranged between 712 and 845; and during the supercolonization experiments, between 773 and 801. Overall, water quality parameters in the tanks were within the optimal ranges for the zebrafish (126). The only substantial environmental difference between the mFTS and the RAS was the temperature. In the RAS, the water temperature is maintained at 28.5 °C by an in-line heating element. In the mFTS, tanks were filled with conditioned RAS water but then equilibrated to air temperature over time. Water temperature in the tanks after the water exchange was recorded daily in the two studies. During the *in vivo* exposure TBT, the average water temperature recorded in the afternoon was 22.3 °C; and in the supercolonization experiments, 22.9°C. Notably, the water temperature equilibrated with the room temperature during the evening, resulting in a lower water temperature in the morning. During the *in vivo* exposure to TBT, the average water temperature recorded in the morning was 19.1°C, and in the supercolonization experiments 20.1°C. Because the mFTS was located inside our main zebrafish facility, we were not able to change the room temperature or install space heaters.

During their housing in the mFTS, zebrafish received an exclusively live feed diet. *Artemia* (Brine Shrimp Direct, BSEACASE) was offered twice a day. *Artemia* was collected, rinsed and resuspended in RAS water. The volume and the concentration of *Artemia* (volume collected from hatching cone/volume of RAS water used for resuspension after rinsing) were adjusted to prevent the accumulation of uneaten food at the bottom of the tanks and recorded daily. Each tank in the mFTS received the same amount of food, and the order of feeding was alternated daily to prevent any feeding bias that could affect growth. Uneaten food and debris accumulated at the bottom of the tank were removed as required using a 10 ml sterile serological pipette during the water exchange.

### Repeated cross-sectional study of in vivo TBT exposure

Four stocks of 1 mg/ml, 0.1 mg/ml, 0.01 mg/ml, and 0.001 mg/ml of TBT (Sigma, T50202) in DMSO (Fisher, D136-L) were prepared. All stocks were maintained in the dark at 4° C until use. Small volumes of these stocks were aliquoted in sterile borosilicate glass vials to avoid repeated freeze-thaw cycles. DMSO stocks were aliquoted in the same manner.

Tanks in the mFTS were initially dosed with 75 µl of the corresponding stocks of TBT and DMSO stocks to achieve final concentrations of TBT in water of 10 μg/l, 1 μg/l, 0.1 μg/l, and 0.01 μg/l, respectively; as well as DMSO vehicle control. By aliquoting the same volume of stocks, we also maintained an equal volume of DMSO in all treatments. Stocks were added before refilling tanks after the water exchange to facilitate mixing. We assumed that TBT was not degraded over a 24-hour period, so we added a volume of stock daily to adjust the concentration of TBT in the tanks for the total loss of chemical during the daily water exchange.

Each TBT concentration and the control had four tank replicates and 30 fish per tank (Fig. 1A). The TBT concentration of 10 µg/l resulted in a mortality rates between 92.31 and 100% after one day of exposure so this treatment was excluded from the study and the schematic in Fig. 1A. The tank replicates for each treatment were randomly distributed in our zebrafish facility racks to avoid feeding and water exchange biases (not represented in Fig. 1A). Four time points were sampled during this repeated cross-sectional study: one before starting the exposure to TBT (0 days post-exposure, dpE); as well as 7, 14, and 21 dpE (Fig. 1A). The fish were not offered food on the day of sampling and had received their last meal and chemical treatments 14 hours before sampling. 200 ml of water from each tank were collected in sterile conical vials: 150 ml were used for the aquatic microbial community analysis (see below), and the remainder 50 ml we used for measuring the water quality parameters (pH, nitrate, nitrite, and toxic ammonia) and for counting colony forming units (CFU). Five individual fish per tank (20 fish per treatment) were sampled at each time point indicated in Fig. 1A. Body size (SL, HAA, and body area), adipose tissue accumulation, and composition of the intestinal microbial communities from each fish were also analyzed (see below, Fig. 1A).

### Zebrafish body size and adipose tissue analysis

Nile Red (Sigma, N1142) staining was performed as described in (33, 70, 78) except that the Nile red solution was prepared with filtered and autoclaved RAS water (FASW). Zebrafish were euthanized in 1.34g/L buffered Tricaine and laid on 1.5% methylcellulose in FASW on the right flank for imaging. We used a Leica M205 fluorescence stereomicroscope with a GFP bandpass filter to image Nile Red fluorescence in adipose tissues and corresponding brightfield images of each animal. All images per time point were acquired using the same exposure settings. Image analysis was conducted in FIJI/ImageJ 1.5p (127). SL, HAA, and total body area were measured to evaluate body size as described in (70, 77, 78).

### Intestinal and aquatic sample collection

After imaging, euthanized zebrafish were transferred to a Petri dish coated with sterile 3% agarose, onto which fish were immobilized by the tail using an insect pin dipped in 70% ethanol. After decapitation with an autoclaved razorblade, the whole gastrointestinal (GI) tract was removed using forceps. Individual whole intestines were separated from the other organs of the GI and transferred to sterile screw cap tube containing 100 µl of sterile 0.1-mm-diameter zirconia beads (BioSpec Products, NC0268065) and 400 µl of sterile lysis buffer (20 mM Tris-HCl pH 8.0, 2 mM EDTA pH 8.0, 1% Triton X-100; modified from (72)). Tank water from the upper water column was filtered as described in (72). Filters were then extracted using flame-sterilized forceps and cut in half using a sterile scalpel. One half of the paper disc was rolled and inserted into a sterile screw cap tube containing 200 µl of sterile 0.1-mm-diameter zirconia beads, to which 200 µl of sterile lysis were immediately added. Intestinal and water samples were flash frozen in a dry ice-ethanol bath and stored at −80°C for subsequent 16S rRNA gene sequencing.

### Supercolonization with zebrafish intestinal bacterial strains

In the supercolonization experiments described here, we used four bacterial strains isolated from the zebrafish intestine: *Chitinibacter* ZOR0017, *Chitinibacter* ZOR0013, *Plesiomonas* ZOR0011; and *Vibrio* ZWU0020 (64, 80). Strains ZOR0017, ZOR0013, and ZOR0011 were generously provided by Karen Guillemin (University of Oregon). Single colonies grown on tryptic soy agar (TSA) plates were picked and grown overnight in tryptic soy broth (TSB). 750 µl of the overnight cultures were collected to extract DNA and confirm the identity of the strain by sequencing the 16S rRNA gene using the universal 8F and 1492R primers. An aliquot of this overnight verified culture was diluted and plated on TSA for growth of well-spread single colonies that were used to inoculate fresh daily overnight cultures used for preparing the bacterial suspension administered daily during the supercolonization experiments. The daily overnight cultures of 10 ml of tryptic soy broth (TSB) were incubated aerobically at 30 °C for 14 to 16 h in rotary mixer at 100 rpm. 300 µl of the daily overnight culture were collected for 16S rRNA sequencing-based verification of the bacterial strains.

A starter culture of 10 ml of fresh TSB was inoculated with the overnight culture at a starting OD_600_ of 0.1 and incubated for 6 to 8 h at 30°C in a roller drum at 100 rpm. The starter culture was used to inoculate 200 ml of fresh sterile TSB in an culture flaks (i.e. supercolonization culture). Because of differences in the growth rate of the different strains, and to obtain supercolonization cultures of approximately the same density at the same time, the supercolonization culture for *Chitinibacter* ZOR0017 and *Chitinibacter* ZOR0013 was inoculated at a starting OD_600_ of 0.05, whereas *Plesiomonas* ZOR0011 and *Vibrio* ZWU0020 were inoculated at a starting OD_600_ of 0.025. The supercolonization culture was incubated at 30°C in a shaker at 100 rpm for 14 hours.

To prepare the bacterial suspension, we calculated the volume of the supercolonization culture required to prepare 200 ml of bacterial suspension in fresh TSB at an OD_600_ of 1. The diluted bacterial suspension was transferred to an autoclaved Nalgene centrifuge bottle (B1033, Sigma), and spun in a Sorvall RC5B centrifuge at 5000rpm at a set temperature range of 10 – 25°C. The pellet was washed twice with 100 ml of autoclaved RAS water, and resuspended in 60 ml of autoclaved RAS water (i.e., bacterial suspension for daily supercolonization). 750 µl of each bacterial suspension were sampled daily to count CFU on TSA and 16S rRNA sequencing-based verification of the bacteria strains. The CFU/ml in the tanks after supercolonization (estimated from the CFU/ml in the bacterial suspension) was measured daily. The average CFU/ml was not significantly different amongst the bacterial strains over the 14 days of supercolonization (One-way ANOVA and Tukey’s multiple comparisons test; *P* = 0.7770). The bacterial suspension was divided into 20 ml aliquots used to supercolonize each tank. Before adding the bacterial suspension, the tanks were drained to the 2.5 L volume. To promote supercolonization of the intestine with this inoculum, zebrafish were offered *Artemia* after adding the bacterial suspension. Zebrafish, bacteria, and *Artemia* were incubated for 2.5 h, before continuing with the water exchange as described above. Control tanks received a mock inoculation of 20 ml of autoclaved RAS.

Each bacterial treatment and the control had three tank replicates (15 experimental tanks total with ∼30 fish per tank) randomized in order. We performed daily supercolonization for 14 days, which was a period equivalent to the earliest time point at which we observed body size and adiposity phenotypes during the *in vivo* chemical exposure (Fig. 2). Five fish per tank were sampled at (0 dpSC) for intestinal microbiota and body size analysis. Fifteen fish were sampled at the 14 dpSC (as there was only one time point in this experiment) for body size and adipose tissue analysis. Five of these fifteen fish were also sampled for intestinal microbiota analysis (Fig. 1B). Tank water was collected from each tank for aquatic microbiota analysis at 0 and 14 dpSC as previously described for aquatic microbiota and water quality analysis.

### Gnotobiotic zebrafish husbandry and obtention of experimental cohorts

The maintenance of the flasks containing GF zebrafish, diet administered and monoassociation procedures were adapted from (108, 128) to maintain GF animals up to 35 dpf. This substantial improvement of gnotobiotic husbandry was necessary for our experiment because adipose tissue starts to develop when zebrafish are SL of ∼4.2 mm which in conventional husbandry is detectable around 12 dpf (69, 70). Moreover, we wanted to mimic the phenotypical *in vivo* chemical exposure to 1 µg/l TBT (both body size and adiposity at 21 dpE; Fig.2) and supercolonization with *Plesiomonas* ZOR0011 (at 14 dpSC; Fig.4) under conventional husbandry conditions.

Four conventionally reared adult fish (1:1 male to female ratio) from two different families were set in mating boxes for group spawning. Zebrafish embryos were obtained by natural mating. The collection of the embryos and the derivation procedure to obtain GF zebrafish was conducted as described in (108). All procedures after GF derivation were conducted using sterile technique, sterile materials wiped with ethanol and UV-irradiated, inside a laminar flow cabinet. Sterility tests throughout this experiment were conducted as described (108). Zebrafish were maintained in cell culture flasks with sterile gnotobiotic zebrafish medium (GZM) in a 28.5°C incubator with a 14:10 hour light: dark cycle. Flasks were filled with 50 ml of GZM from 0 to 21 dpf, and this volume was increased to 100 ml GZM from 23 to 35 dpf to accommodate the growth of the animals. 60% volume GZM exchange was conducted daily to maintain water quality levels. pH, nitrate, nitrite, and toxic ammonia were measured in all flasks as previously described. The pH in the flasks ranged between 6.6 and 7, nitrite between 0 and 0.15 ppm, nitrate between 0 and 10 ppm, and toxic ammonia between 0 and 0.0042 mg/l. The pH in the flasks was lower than in the conditioned water in our RAS facility and the *in vivo* chemical exposure, but comparable to the pH during the supercolonization experiments. The nitrite in the flasks was higher than in the conditioned water in our RAS zebrafish facility and the supercolonization experiments, but comparable to the *in vivo* chemical exposure. The nitrate in the flasks was comparable to the conditioned water in our RAS zebrafish facility, the *in vivo* chemical exposure and the supercolonization experiments. The toxic ammonia in the flasks was higher than in our RAS zebrafish facility, but comparable to the *in vivo* chemical exposure and the supercolonization experiments. Overall, water quality parameters in the flasks were within the optimal ranges for zebrafish (126).

Flasks of GF zebrafish were raised for 13 days post-fertilization before randomly assigning flask to the chemical and microbial treatments (see below). These flasks also underwent a thorough fish health check and verification of GF status. On 14 dpf, one fish per flask was sampled to measure the SL. The average SL of the cohort was 4.21 ± 0.22 mm (n=29), similar to the SL expected at this age under conventional husbandry. Exposure of GF zebrafish to TBT and their monoassociation with *Plesiomonas* ZOR0011 or *Chitinibacter* ZOR0013 were initiated on 15 dpf. Flasks of GF zebrafish (5 flasks per treatment except for *Chitinibacter* ZOR0013 which had only four, 19 flasks total) were randomly assigned to chemical and bacterial treatments. Two time points were sampled: 7 and 21 days post-exposure (dpE) or monoassociation (days post-colonization, dpC). Flasks were sampled terminally at both time points: two flasks per treatment at 7 dpE, and 3 samples per treatment at 21 dpE (except for the *Chitinibacter* ZOR0013 treatment from which only two flasks) were sampled at each time point. The fish were not offered food on the day of sampling and had received their last meal and treatments 14 hours before sampling. Body size and adipose tissue area from each fish in the flask were analyzed as previously described. There were no significant differences in the rearing density of the flasks assigned to the chemical/bacterial treatments that could result in differences in body size (Kruskal-Wallis, Dunn’s multiple comparison test of flasks from each chemical/bacterial treatment against the germ-free DMSO control; all *P* ≥0.8869 at 7 dpE/dpC; and *P* ≥ 0.9474 at 21 dpE/dpC). Similarly, there were no differences in mortality in flasks sampled at 7 or 14 dpE/dpC (Kruskal-Wallis, Dunn’s multiple comparison test of flasks from each chemical/bacterial treatment against the germ-free DMSO; all *P* ≥0.1986 at 7 dpE/dpC; and *P* ≥ 0.5213 at 21 dpE/dpC).

### Culture and provision of germ-free Artemia-Tetrahymena diet

One of the main challenges of long-term gnotobiotic husbandry is the transition in zebrafish larvae from reliance on yolk to free-feeding before they are capable of hunting live *Artemia*. Also, we wanted to reproduce as closely as possible the diet zebrafish received in the mFTS during the *in vivo* chemical exposure, and the supercolonization experiments. However, previous pilot experiments showed that an *Artemia*-only diet was not compatible with survival past 14 dpf (data not shown). Thus, we designed a GF live diet of *Tetrahymena thermophila* strain CU428.2 (TSC_SD00178, Tetrahymena Stock Center) and San Francisco strain *Artemia* (Brine Shrimp Direct). GF *Artemia* was offered daily starting 5 dpf, and supplemented with GF *T. thermophila* on 5, 8, and 11 dpf. GF *Artemia* and *T. thermophila* cultures were prepared as described below.

*T. thermophila* stocks were maintained in 5 ml of Modified Neff’s Media (MNM; 3 mg FeCl_3_.6H_2_0, 10 g proteose peptone, 2.5 g yeast extract, and 5 g glucose) with 5 μl Penicillin-Streptomycin Stabilized solution (Sigma-Aldrich, P4458) and 24 μl of Amphotericin B solution (Sigma-Aldrich, A2942) at room temperature. These stocks were renewed every two weeks by transferring 100 µl of stock to fresh media. 200 µl of MNM stock were used to inoculate 14.8 ml of Proteose Peptone Yeast Agar (PPYE; 2,5 g proteose peptone; 2.5 g yeast extract) media with 10 µl of Penicillin-Streptomycin Stabilized solution (Sigma-Aldrich, P4458). PPYE cultures were maintained at 28.5° C for one week before its use for inoculation of milk yeast extract media (MYE; 10g/l yeast extract, 10 g/l non-fat powdered milk). MYE is a rich media that is used for growing *T. thermophila* for feeding zebrafish (129). 4 ml of a one-week old PPYE culture were used to inoculate 40 ml of sterile MYE and was incubated at 28.5° C overnight in a 50 ml conical vial. For food preparation, the conical vial was centrifuged at 5000 rcf for 3 minutes, 25 ml of the supernatant were decanted, and the conical vial was refilled to the 50 ml mark with sterile GZM to rinse the pellet. The pellet was rinsed two more times with sterile GZM by decanting 35 ml of the supernatant, before its final resuspension in 40 ml of sterile GZM. This preparation had an average of ∼30 000 cells/ml. 4 ml of the suspension of *T. thermophila* were used to feed each fish flask. Sterility tests were conducted on the *T. thermophila* MNM and PPYE cultures, as well as on prepared food suspensions as described in (108).

To prepare GF Artemia, ∼250 µl of Artemia cysts were rehydrated in 10 ml of sterile saltwater (12.5g/l NaCl, 0.83g/l NaHCO3) for at least 2 hours with gentle shaking in a nutator. The cysts were left to settle, and the saltwater was exchanged for 10 ml of household bleach (8.25% sodium hypochlorite) for decapsulation. The Artemia-bleach suspension was mixed on a nutator for 3 minutes, and then left to settle at the bottom of the conical tube for 5 more minutes. After decapsulation, the bleach and floating cysts were removed. The cysts were rinsed by adding saltwater, mixed by gentle rolling for 1.5 minutes, before letting the cysts settle for 30 seconds. The procedure was repeated 4 times, before resuspending the cysts in 10 ml of sterile saltwater, and incubation for at least 24 h at room temperature with gentle shaking under a warm light for hatching. For food preparation, the actively swimming *Artemia* were collected using a sterile serological pipette, collected on a 100 µm cell strainer (431752, Corning) and rinsed with 10 ml of sterile GZM. *Artemia* was transferred to a conical vial and resuspended in GZM. The volume and the concentration (i.e. volume of collected from hatching vial/volume of GZM for resuspension) of *Artemia* was adjusted to prevent the accumulation of food debris in the flasks. Volume and concentration were recorded daily. Each flask of fish received the same amount of food, and the order of feeding was alternated daily to prevent to prevent any feeding bias that could affect growth. Uneaten food and debris were removed daily during the GZM exchange.

### Germ-free zebrafish exposure to TBT

Stocks of 1 mg/ml of TBT (Sigma, T50202) in DMSO (Fisher, D136-L) and of DMSO vehicle were prepared and aliquoted using sterile technique. Although the presence of bacteria in these chemicals was unlikely, the sterility of the stocks was tested by plating 100 µl of the stock solutions on TSA and incubated for 1 week at 28.5°C under aerobic conditions before starting the experiment. The plates were inspected daily, and no microbial growth was observed.

Flasks containing GF zebrafish were initially dosed with 5 µl of the TBT stock and DMSO to achieve a final concentration of 1 μg/l TBT as well as an equal total volume of DMSO vehicle in the control flasks. Stocks were added before fresh GZM during the media exchange to facilitate mixing. We assumed that TBT was not degraded over a 24-hour period, so we added a volume of stock daily to adjust the concentration of TBT in the flask for the total loss of chemical during the daily GZM exchange.

### Monoassociation with Plesiomonas ZOR0011 and Chitinibacter ZOR0013

Monoassociation is the colonization of GF zebrafish with a single microbial strain. A starter culture of each bacterial strain was prepared by inoculating 10 ml of sterile TSB with a single colony, followed by overnight aerobic incubation in a roller drum at 100 rpm at 30° C. 300 µl of the starter culture were collected to confirm the identity of the strain by sequencing as described for the supercolonization experiments. The starter culture was used to inoculate 40 ml of sterile TSB in a sterile 50 ml conical via (i.e. monoassociation culture). To obtain monoassociation cultures of approximately the same density at the same time, *Chitinibacter* ZOR0013 was inoculated at a starting OD_600_ of 0.05 and *Plesiomonas* ZOR0011 at 0.025 as *Plesiomonas* ZOR0011 grew faster in TSB than *Chitinibacter* ZOR00R13. The monoassociation culture was incubated at 30° C in a shaker at 100 rpm for 14 hours. To prepare the bacterial suspension for the monoassociation, we calculated the volume of the culture needed to prepare 40 ml of a bacterial suspension at an OD_600_ of 0.25. We chose the OD_600_ and dilutions to ensure that the zebrafish fish received amounts of bacteria in the monoassociation comparable to those in the supercolonization experiments. The diluted bacterial suspension was spun at 5000 rpm for 10 minutes. The supernatant was decanted, and the pellet was washed twice with 40 ml of sterile GZM. The washed bacterial pellet was finally resuspended in 40 ml of autoclaved GZM, and this suspension was used for the monoassociation. 750 µl of each bacterial suspension were sampled to count CFU on TSA plates and to confirm the identity of the strain by 16S rRNA gene sequencing as described above. The CFU/ml in the flask after inoculation were of 1.01E+08 CFU/ml for *Chitinibacter* ZOR0013 and 1.20E+07 CFU/ml for *Plesiomonas* ZOR0011 (estimated from the CFU/ml of the bacterial suspension). Nevertheless, the average CFU/ml counts from GZM from individual flasks sampled at 7 dpC and 21 dpC were not significantly different (t-test, *P* = 0.7541 and *P* = 0652 respectively) between *Chitinibacter* ZOR0013 and *Plesiomonas* ZOR0011, suggesting that this initial difference was not maintained and likely did not affect the results of our experiments. In addition, whereas the estimated CFU/ml in the fish flasks after monoassociation was lower than the average estimated CFU/ml in the tanks after the supercolonization experiments, these values were within the minimum to maximum CFU/ml range in tanks after supercolonization measured during the 14 days of supercolonization (data not shown).

GF flasks assigned to the bacterial treatments received 6 ml *Chitinibacter* ZOR0013 or *Plesiomonas* ZOR0011 suspension. In addition, the monoassociated flasks were initially dosed with 5 µl of DMSO since the GF DMSO control was shared with the chemical exposure to TBT in the GF background. DMSO was added before fresh GZM during the media exchange to facilitate mixing, and a volume was added daily to adjust for the loss chemical during the daily GZM exchange. Bacteria were added only once during the experiment at 15 dpf.

### In vitro exposure of zebrafish intestinal microbial communities to TBT

Four stocks of 1000 µg/ml, 10 µg/ml, 1 µg/ml, 0.1 µg/ml, and 0.01 µg/ml of TBT (Sigma, T50202) in DMSO (Fisher, D136-L) were prepared and aliquoted using sterile technique. All stocks were maintained in the dark at 4°C until use. Borosilicate tubes containing 15 ml of sterile tryptic soy broth (TSB) were used for the aerobic *in vitro* exposure; and borosilicate tubes containing 14.25 ml of sterile TSB and 0.75 µl of 10 g/l filter sterilized L-cysteine were used for the anaerobic *in vitro* chemical exposure. 15 µl of the stocks were added to the growth media to achieve final TBT concentrations of 0 µg/l (DMSO vehicle control), 0.01 μg/l, 0.1 μg/l, 1 μg/l, and 10 μg/l, and 1000 μg/l TBT. Each TBT concentration was prepared in triplicate.

An experimental cohort of zebrafish was obtained and conventionally reared. On day 25 post-fertilization, zebrafish were size matched between 7 mm and 9 mm. Two pools of 30 size-matched individuals were transferred to two clean and autoclaved fish tanks and maintained in the RAS for 4 weeks before sampling to match the SL at 21 dpE *in vivo* chemical exposure to TBT. Two fish per tank were collected as donors (n = 4; average SL = 16.75 ± 1.5 mm). To reproduce the sampling of whole intestines during the *in vivo* chemical exposure, zebrafish were incubated in 1 mM epinephrine in FASW before euthanasia with 1.34 g/L filter sterilized tricaine. Whole gastrointestinal (GI) tracts were dissected as previously described, and cut open using sterile technique before transfer to a sterile screw cap tube containing 200 µl of sterile 1-mm-diameter zirconia beads (NC9847287) and 500 µl of FASW. Four GI tracts were pooled in a single beaded screw cap tube. Intestinal samples were vortexed for one minute twice to obtain an intestinal slurry used to inoculate TSB media with TBT and DMSO prepared as described above (10 µl of slurry in 5 ml of media). The remainder of the slurry constituted the input intestinal microbial community and was transferred to a sterile Eppendorf tube. The beads were rinsed with 500 µl of FASW, and after vortexing, this rinsing volume was pooled with the slurry. The collected material was centrifuged, before centrifugation at 7500 rcf for 15 minutes. After removing the supernatant, the pellet was flash frozen in a dry ice-ethanol bath and stored at −80°C until DNA isolation for 16S rRNA gene sequencing.

TSB cultures dosed with TBT or DMSO were incubated aerobically and anaerobically for 24 hours without agitation at room temperature. Anaerobic incubation was performed in a Coy anaerobic chamber. 750 µl of each culture tube was collected in a sterile 1.5 ml Eppendorf tube and spun at 7500 rcf. The pellets were flash frozen in a dry ice-ethanol bath and stored at −80°C until DNA isolation for 16S rRNA gene sequencing for microbial community analysis.

### DNA isolation

The Duke Microbiome Shared Resource extracted bacterial DNA from all samples using Qiagen MO BIO PowerSoil DNA Kit that allows for the isolation of samples in a 96 well plate format using the Retsch MM400 plate shaker. Quantity of samples were assessed using the PerkinElmer Victor plate reader with the ThermoFisher Scientific Qubit dsDNA HS assay kit.

### Sequencing of 16S rRNA gene amplicons

Bacterial community composition in isolated DNA samples was characterized by amplification of the V4 variable region of the 16S rRNA gene by polymerase chain reaction using the forward primer 515 and reverse primer 806 16S rRNA following the Earth Microbiome Project protocol (http://www.earthmicrobiome.org/) by the Duke Microbiome Shared Resource. These primers (515F and 806R) carry unique barcodes allowing for multiplexing and co-sequencing of hundreds of samples at a time. Equimolar 16S rRNA gene PCR products were quantified and pooled for sequencing. Sequencing was performed by the Duke Microbiome Center using an Illumina MiSeq instrument configured for 150 bp paired-end sequencing runs.

### Bioinformatic analysis

Data analysis conducted in QIIME v. 1.9.1 (131). Paired reads were joined with join_paired_ends.py, using standard parameters, and joined reads were subsequently demultiplexed with split_libraries_fastq.py, with the phred quality threshold set to Q30. Demultiplexed amplicons were filtered for chimeric sequences with identify_chimeric_seqs.py, using the usearch61 (132) detection method, and substituting usearch v. 6.1 with vsearch v. 2.7.1 (133). Only amplicons predicted as chimeric by both denovo and reference-based detection were removed. After completion of base quality and chimera filtering, totals of 21,069,394 (*in vivo* TBT exposure study), 1,283,416 (supercolonization study) and 5,801,110 (*in vitro* TBT exposure study) amplicons were used in downstream analyses. Amplicons were clustered into operational taxonomic units (OTUs), using QIIME’s closed reference OTU picking strategy, implemented in pick_closed_reference_otus.py. The script was configured to cluster amplicons with uclust (132) and assign taxonomy against the 97% clustered 16S reference sequence set of SILVA, v. 1.23 (134). A subsequent basic statistical diversity analysis was performed, using QIIME’s core diversity analysis workflow script (core_diversity_analyses.py), calculating alpha- and beta-diversity, and relative taxa abundances in sample groups. Relative taxa abundances were further analyzed with LEfSe (Linear discriminant analysis effect size, (79) to identify differential biomarkers in sample groups.

### Data availability

16S rRNA sequence data are available at the NCBI Short Read Archive under accessions PRJNA692784 (*in vivo* TBT exposure study), PRJNA693006 (supercolonization study), PRJNA693015 (*in vitro* TBT exposure study).

## ACKNOWLEDGEMENTS

This work was supported by grants to J.F.R. from the National Institutes of Health (R21-ES023369) and the Gordon and Betty Moore Foundation. The authors are grateful to Karen Guillemin and Brendan Bohannan (University of Oregon) for providing bacterial strains; José M. Abelleira-Pereira (Universidad de Cádiz) for assistance in design of the mFTS; Philip Zandona for technical assistance in construction of the mFTS; Jim Burris and Patrick Williams (Duke Zebrafish Core Facility) for accommodation and support of our zebrafish experiments; Holly Dressman and Zhengzheng Wei for their assistance in the optimization of extraction protocols and other helpful conversations; Colin Lickwar and Jia Wen for helpful feedback on this manuscript; and the competent staff at Public Hardware Inc. (Durham, NC).

## Notes

### Competing Interest Statement

The authors have declared no competing interest.

### Summary of Updates

Supplemental figures and tables added.

