## Supplemental Figures 1 and 2 for "Tributyltin exposure leads to increased adiposity and reduced abundance of leptogenic bacteria in the zebrafish intestine"

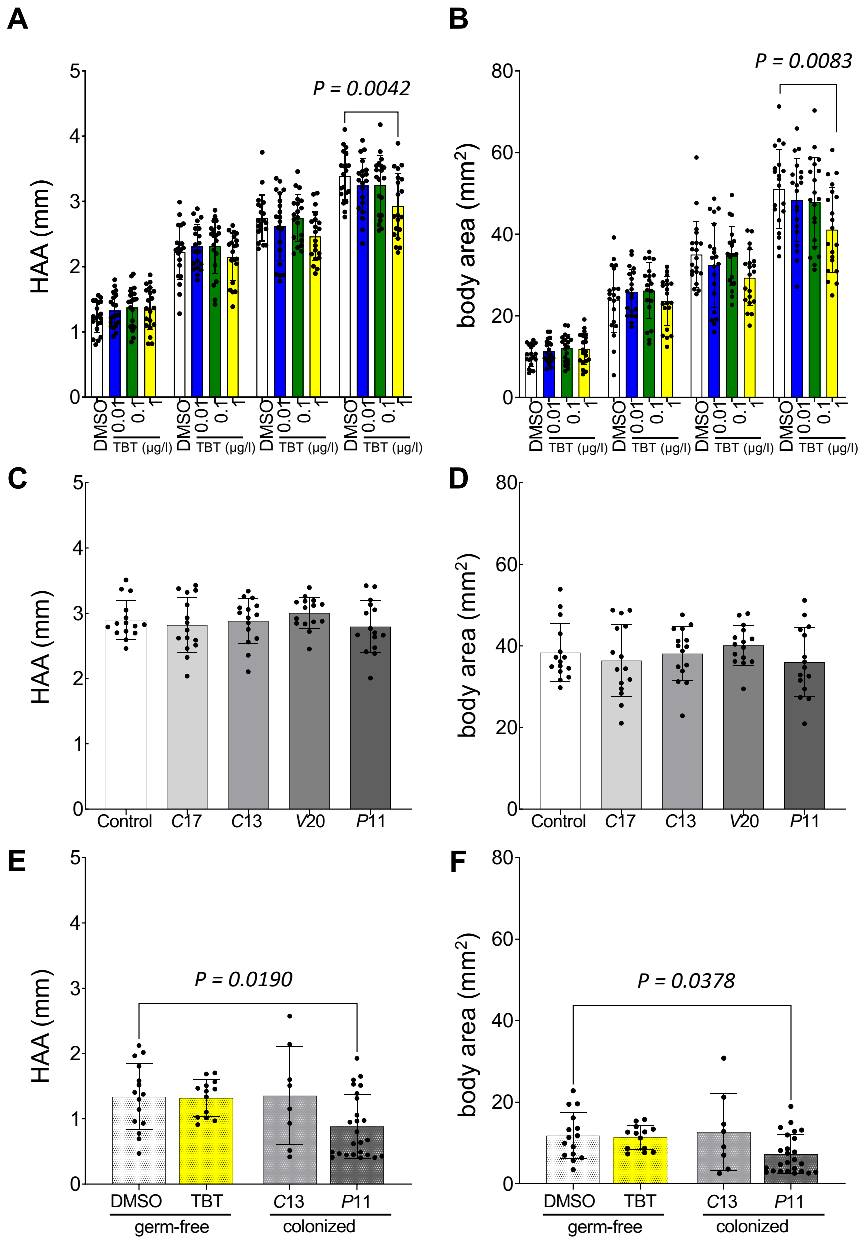


**Figure S1: Effect of bacterial and chemical treatments on additional measurements of body size.** (A) Height at anterior of anal fin, HAA and (B) body area during the exposure to TBT. (C) HAA and (D) body area after 14 days of supercolonization with four bacterial strains: *Chitinibacter* ZOR0017, *C*17; *Chitinibacter*

ZOR0013, *C*13; *Vibrio* ZWU0020, *V*20; or *Plesiomonas* ZOR0011, *P*11. (E) HAA and (F) body area after 14 days of exposure of germ-free zebrafish to TBT; or monoassociation (days post-colonization, dpC) with *Chitinibacter* ZOR0013 (*C*13) and *Plesiomonas* ZOR0011 (*P*11). Bars represent the mean and error bars the standard deviation. Means of the different treatments were compared to their respective controls using one-way ANOVA and post-hoc Dunnett's multiple comparison tests. The controls were: DMSO in (A) and (B); recirculating aquaculture system water in (C) and (D); and germ-free DMSO in (E) and (F). When significant, the multiplicity adjusted *P*-value is reported for the comparison.

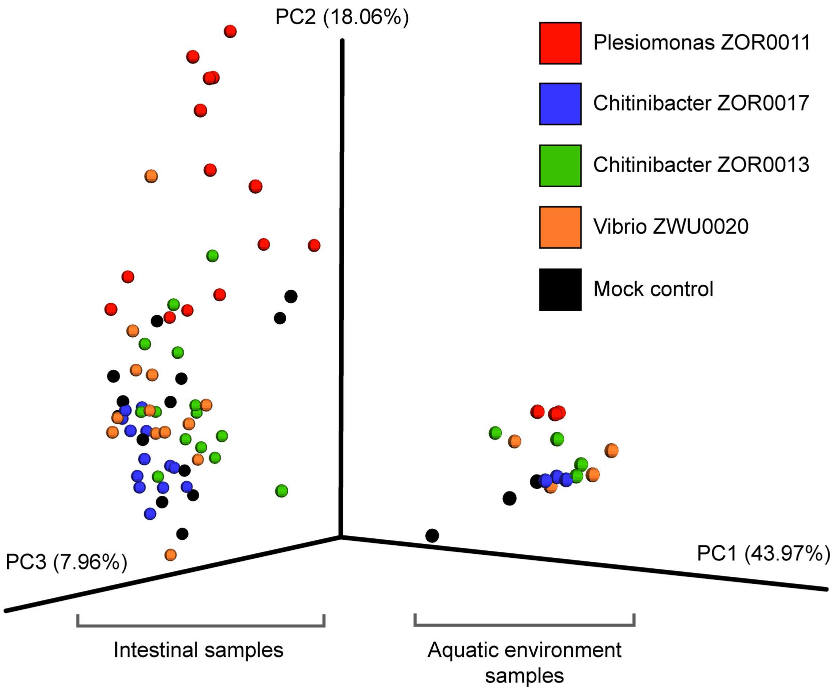


**Figure S2: Impact of supercolonization on microbiota composition in zebrafish intestine and aquatic environment.** Principal coordinates analysis (PCoA) plot based on Bray-Curtis dissimilarity index derived from 16S rRNA gene sequences from intestinal and aquatic environment samples from the supercolonization experiment. The percent variation explained by each of the three PC vectors is shown. Each replicate sample is represented by a single sphere. Note the major differences between intestinal and aquatic environment samples, and the impact of Plesiomonas ZOR0011 supercolonization on overall intestinal microbiota composition. See also Table S4 and S5 for biomarker taxa linked to these treatments, and Table S6 for the PERMANOVA analysis results.
